# BryoCAD: Extending Plant DNA Foundation Models Beyond Flowering Plants

**DOI:** 10.64898/2026.11.09.765485

**Authors:** Ming-Ren Yen, Jingjing Zhai, Sheng-Kai Hsu, Ting-Ying Wu

## Abstract

Bryophytes exhibit diverse genome architectures and gene repertoires, yet their functional interpretation remains constrained by incomplete genomic annotations and limited experimental resources. Although DNA foundation models offer promising solutions, their applicability to evolutionarily divergent bryophytes remains largely unexplored. Here, we introduce BryoCAD, a DNA language model developed through continued pre-training of the angiosperm-specific PlantCAD2 model on 17 bryophyte genomes spanning hornworts, liverworts, and mosses. Across three representative bryophyte genomes and four gene-boundary motif recovery tasks, BryoCAD-Small improved mean zero-shot accuracy from 45.7% to 57.0%, a 24.8% relative increase over PlantCAD2-Small, while the over 200-fold larger Evo2-20B achieved a mean accuracy of only 39.3%. Remarkably, BryoCAD-Small outperformed PlantCAD2-Large in 11 of 12 motif-recovery comparisons across the three bryophytes, highlighting the benefits of lineage-specific adaptation over model scaling alone. These improvements extended to all six tested algal species, increasing mean accuracy from 26.3% to 37.5%, although performance varied across ferns and gymnosperms. BryoCAD also enhanced gene structure annotation: replacing PlantCAD2-Small with BryoCAD-Small increased transcript-level F1 scores from 0.578 to 0.625 in *Marchantia polymorpha* and from 0.485 to 0.530 in *Physcomitrium patens*. Together, our findings demonstrate that lineage- specific continued pre-training enhances genomic sequence interpretation and gene annotation across bryophytes, providing a transferable framework for extending DNA foundation models to evolutionarily diverse, understudied plant lineages.

## Introduction

Bryophytes comprise approximately 22,000 species worldwide and occupy diverse habitats, including extreme environments^1,2^. They encompass three major lineages, hornworts, liverworts, and mosses, with mosses representing the most species-rich group^3–6^. Phylogenetic evidence supports bryophyte monophyly and their sister relationship to vascular plants (tracheophytes), indicating that these two major land plant lineages diverged about 500 million years ago^7,8^. This deep evolutionary divergence has likely contributed to distinct patterns of genome evolution and diversification. Indeed, a recent comparative analysis of 123 bryophyte genomes revealed substantially greater gene family diversity than in vascular plants, characterized by an abundance of unique and lineage-specific gene families and extensive gene family innovation^3^. These findings suggest that bryophytes have evolved distinct genomic strategies that may contribute to their adaptation across diverse terrestrial environments. Given their evolutionary divergence and unique genomic features, analytical tools developed primarily for vascular plants may not fully capture bryophyte genomic complexity, highlighting the need for lineage-adapted computational approaches.

In recent years, rapid advances in genomic language models (LMs) have helped bridge the gap between genomic sequences and functional annotations by learning sequence features directly from DNA and transferring knowledge across species^9–15^. For example, DNABERT, an early DNA language model pre-trained on the human genome, demonstrated its ability to predict regulatory elements^9^. More recently, Evo2 was trained on approximately 9 trillion nucleotides spanning all domains of life, enabling the prediction of variant effects and diverse genomic features^12^. Similarly, Nucleotide Transformer v3 (NTv3) was developed as a multispecies foundation model integrating sequence representation learning, functional genomic prediction, and genome annotation^10^. Among plant-specific models, AgroNT was pre-trained on 48 plant genomes, primarily focusing on crops, but lacked single-nucleotide resolution^11^. Building on its predecessor, PlantCaduceus (PlantCAD), PlantCAD2 introduced an improved Mamba2 architecture, expanded pre-training to 65 angiosperm genomes, and extended the sequence context window to 8,192 bp, achieving improved performance in genomic feature prediction and sequence interpretation across flowering plants^13,15^. Despite these advances, existing DNA foundation models have largely adopted either broad multispecies training strategies or lineage-specific approaches focused on angiosperms, leaving bryophytes underrepresented. Given the importance of evolutionary diversity in learning conserved and lineage-specific genomic features, expanding phylogenetic representation during model pre-training may improve predictive performance in evolutionarily divergent plant lineages.

In addition to predicting genomic features and regulatory elements, accurate gene annotation remains a major challenge, particularly for wild and rare plant species with limited transcriptomic and proteomic resources. This is partly because widely used annotation pipelines, such as BRAKER3, rely on approaches that integrate transcriptomic and proteomic evidence^16^. To reduce dependence on experimental data, GeneCAD was developed by integrating single-nucleotide-resolution representations from PlantCAD2 with a ModernBERT encoder and a chromosome-scale conditional random field that enforces gene-structure constraints. This framework improved transcript-level F1 scores by 8–10% compared with existing annotation methods in angiosperms^17^, highlighting its potential for extension to genome annotation in other plant lineages.

Building on these advances, we developed BryoCAD through continued pre-training of PlantCAD2 on 17 phylogenetically diverse bryophyte genomes. We evaluated its performance in zero-shot motif recovery, variant-effect prediction, chromatin accessibility prediction, and gene structure annotation. We further adapted the GeneCAD framework to improve genome annotation in bryophytes. Notably, BryoCAD also demonstrated improved predictive performance beyond bryophytes, particularly in streptophyte algae. Together, our results establish lineage-specific continued pretraining as a scalable strategy for extending DNA foundation models to evolutionarily divergent plants with limited functional genomic resources.

## Results

### BryoCAD extends plant DNA foundation modeling to evolutionarily diverse bryophytes

PlantCAD2 is a bidirectional, reverse-complement-equivariant DNA foundation model based on the Mamba2 state-space architecture. The model processes 8,192-bp DNA sequences at single-nucleotide resolution and employs a masked language modeling objective to learn sequence-context dependencies^15^. We developed BryoCAD through continued pre-training of PlantCAD2 on 17 phylogenetically diverse bryophyte genomes representing 17 orders across the three major bryophyte lineages, thereby adapting the model to capture bryophyte-specific genomic sequence features (Figure 1A). During continued pre-training, training loss decreased from 1.158 to 1.040 for BryoCAD-S and from 1.136 to 0.927 for BryoCAD-L, corresponding to reductions of 10.2% and 18.4%. respectively. We subsequently designed a series of tasks to evaluate model performance and applied BryoCAD to predict genomic features and improve gene structure annotation (Figures 1B and 1C).

**Figure 1.**
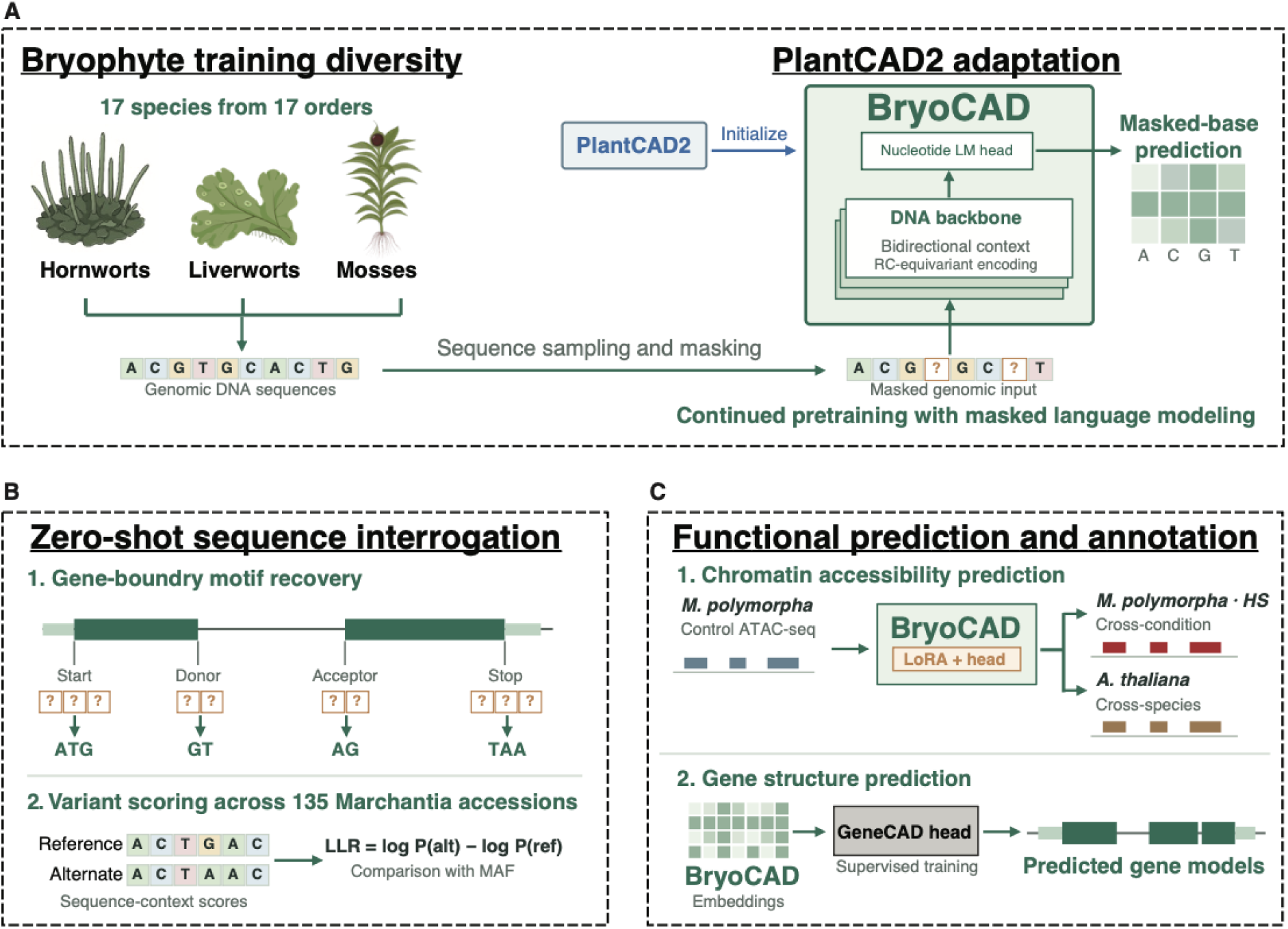
Overview of the BryoCAD framework and downstream applications. (A) Schematic of the continued pre-training strategy used to develop BryoCAD. Genomic sequences from 17 bryophyte species spanning 17 orders were used for continued pre-training. The model architecture, DNA sequence masking strategy, and masked-nucleotide prediction objective were adapted from PlantCAD2. (B) Overview of the zero-shot prediction framework for gene-boundary motif recovery and variant-effect scoring across 135 *Marchantia* genomes. (C) Schematic of the downstream applications of BryoCAD for functional prediction and improved genome annotation. Detailed information on genome sources and species used for pre-training is provided in Supporting Dataset S8.

### Lineage-specific continued pretraining substantially improves motif recovery without increasing model size

We first benchmarked BryoCAD-S (88M parameters) and BryoCAD-L (694M parameters) against PlantCAD2-S (88M), PlantCAD2-L (694M), and three Evo2 models (1B, 7B, and 20B parameters) across seven plant species, including three representative bryophytes and four angiosperms (Figure 2, Supporting Dataset S1). BryoCAD generally outperformed Evo2 across the evaluated tasks and lineages, with the greatest improvements observed in bryophytes. Notably, BryoCAD-S increased mean exact motif recovery accuracy from 45.7% to 57.0% compared with PlantCAD2-S across the three bryophytes, representing an 11.3-percentage-point increase (24.8% relative improvement). In contrast, BryoCAD exhibited only slightly lower prediction accuracy than PlantCAD2 in angiosperms. For example, in angiosperms, BryoCAD-S showed an average decrease of 6.9 percentage points relative to PlantCAD2-S across the four tasks, with the largest relative reduction observed for TTS recovery (26.8%). In addition, we assessed the computational cost of zero-shot motif recovery by benchmarking all seven models on a single NVIDIA RTX PRO 6000 Blackwell GPU using 1,000 sites, equally divided between 2- and 3-bp motifs. BryoCAD-S and BryoCAD-L required an average of 42.03 and 223.22 s per 1,000 sites, respectively, closely matching the inference times of PlantCAD2-S and PlantCAD2-L (41.88 and 223.11 s, respectively). In contrast, Evo2-1B, Evo2-7B, and Evo2-20B required 359.57, 1,246.76, and 2,710.54 s, respectively (Supporting Dataset S1). Together, these results demonstrate that lineage-specific continued pre-training substantially improves motif recovery in bryophytes while largely preserving predictive performance in flowering plants and the inference efficiency of PlantCAD2, even with a relatively small model.

**Figure 2.**
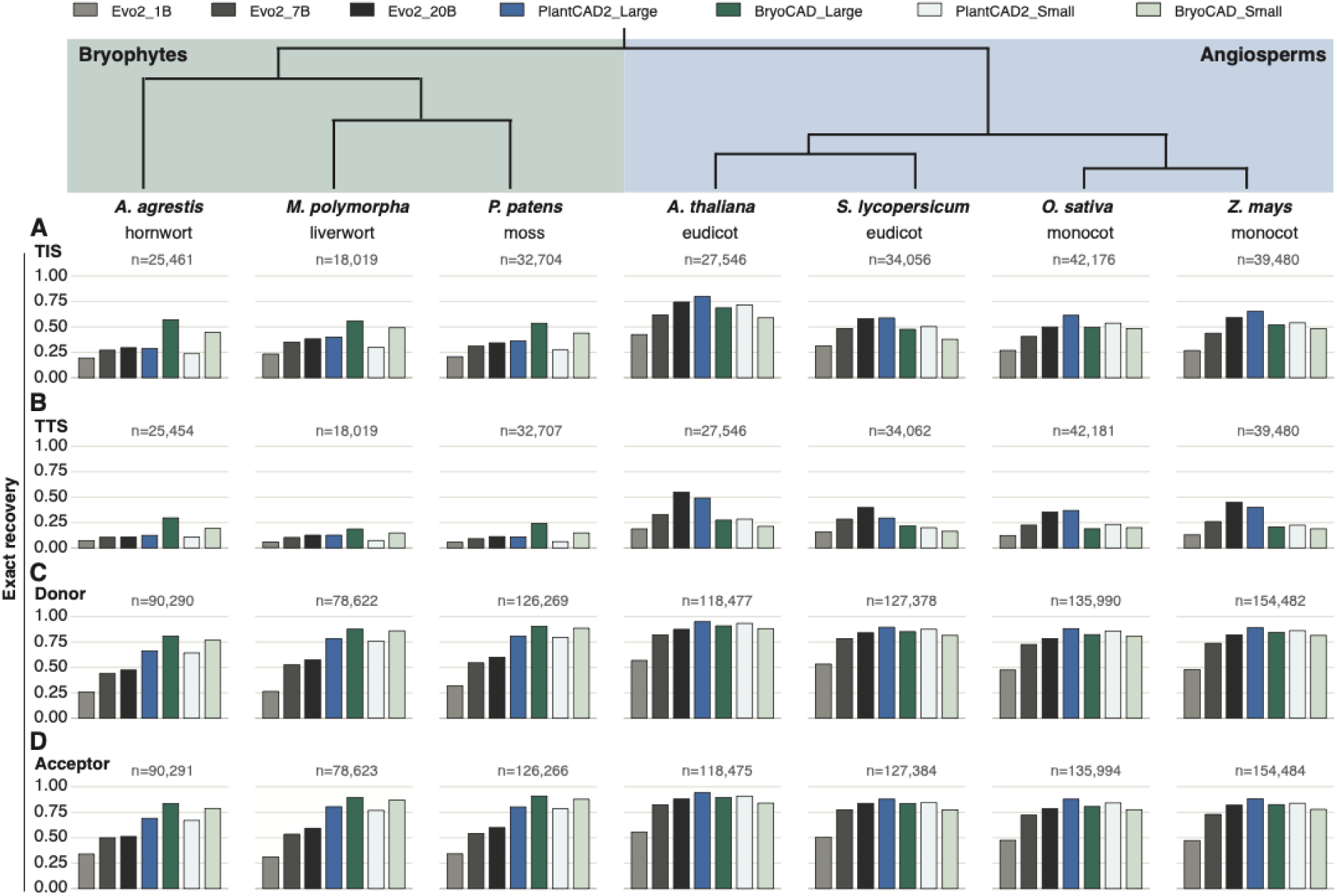
Benchmarking BryoCAD against existing DNA foundation models across seven species. (A–D) Bar plots showing exact motif recovery accuracy across seven species for four genomic boundary prediction tasks: (A) translation initiation sites (TISs), (B) translation termination sites (TTSs), (C) splice donor sites, and (D) splice acceptor sites. Bars are colored by DNA language model, including Evo2 with 1 billion (1B), 7 billion (7B), and 20 billion (20B) parameters; PlantCAD2-S (88 million [88M] parameters) and PlantCAD2-L (694 million [694M] parameters); and BryoCAD-S (88M parameters) and BryoCAD-L (694M parameters).

### BryoCAD generalizes across 126 bryophyte genomes, with prediction accuracy strongly associated with gene conservation

We next extended the evaluation to 126 bryophyte genomes, comprising 5 hornworts, 38 liverworts, and 83 mosses. To assess cross-species generalization, we compared exact motif recovery accuracy among species included in pre-training, unseen species from orders represented during pre-training, and species from orders absent during pre-training. No significant differences were observed among the three groups across all four tasks (Figure 3A; Supporting Dataset S2), suggesting that BryoCAD captures conserved sequence features that generalize across evolutionarily diverse bryophytes. We further examined whether prediction accuracy varied among gene sets with different degrees of conservation. Across all four tasks, mean recovery accuracy was approximately threefold higher for core genes than for unique genes (Figure 3B; Supporting Figure S1, Supporting Dataset S2), suggesting that lineage-specific genes contain less conserved or more divergent sequence features. Finally, we compared model performance across the three major bryophyte lineages. Mosses consistently exhibited slightly but significantly higher recovery accuracy than liverworts across all four tasks (adjusted q < 0.05; Figure 3C). This difference was not clearly explained by genome properties such as genome size, gene density, or GC content (Supporting Figures S3–S5, Supporting Dataset S2), but may partially reflect the higher proportion of unique genes in liverwort genomes (Supporting Figure S1).

**Figure 3.**
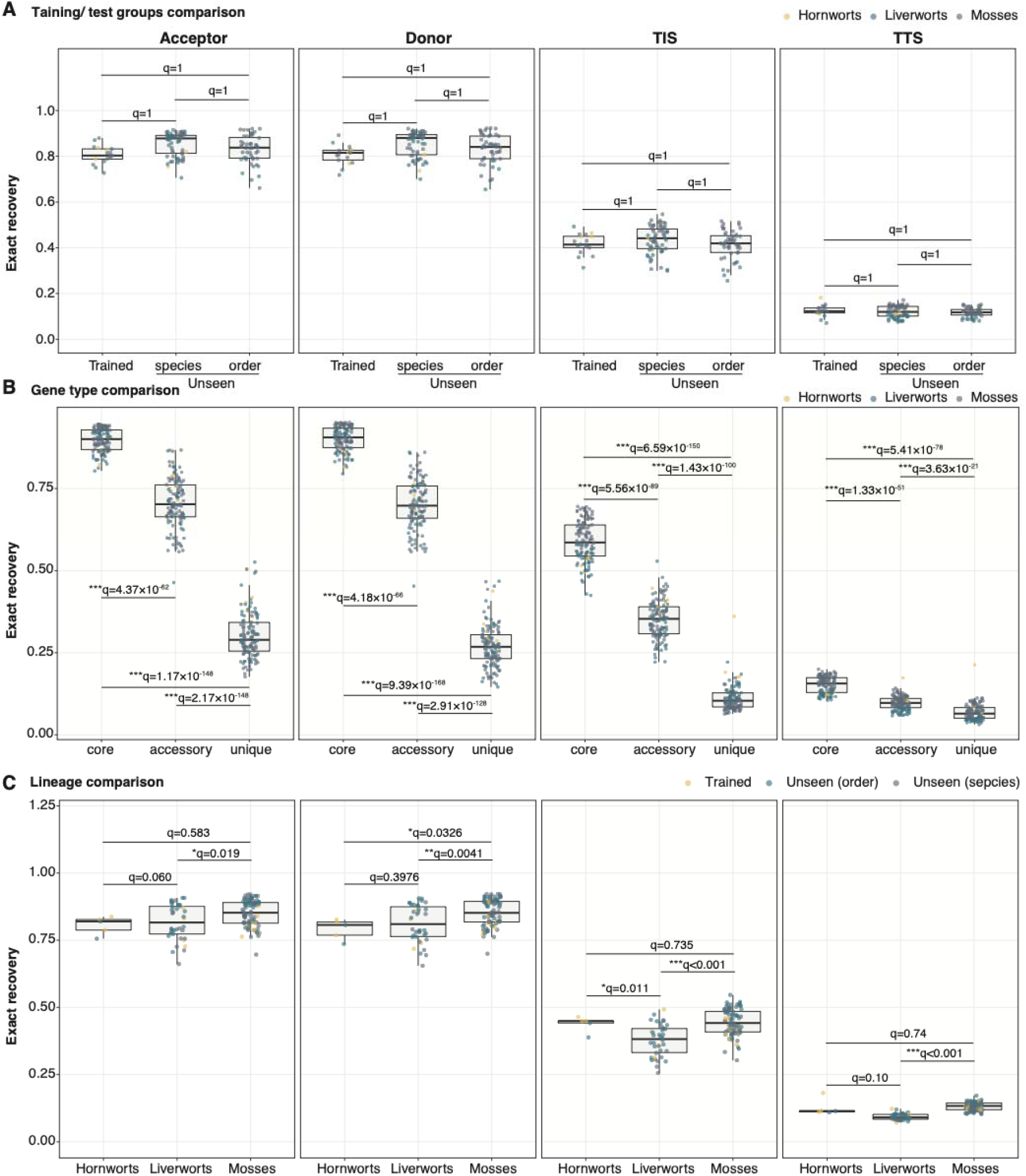
Cross-species generalization of BryoCAD across 126 bryophyte species. (A) Box plots comparing exact motif recovery accuracy among three categories. Seen genomes (n = 17): Assemblies included in training. Unseen species (n = 61): Assemblies excluded from training but belonging to orders represented in training. Unseen orders (n = 48): Assemblies from orders absent from training. (B) Box plots showing exact motif recovery accuracy across different gene sets. Core sets were defined as those present in more than 80% of genomes (101–126 genomes), accessory sets as those present in 2–100 genomes, and unique sets as those restricted to a single genome. (C) Box plots comparing exact motif recovery accuracy among the three major bryophyte lineages: hornworts, liverworts, and mosses. In all panels, dots represent individual data points, center lines indicate medians, and boxes span the first and third quartiles. Statistical significance was assessed using one-sided Welch’s *t*-tests with Benjamini–Hochberg correction for multiple comparisons. Adjusted *P* values are reported as *q* values.

Interestingly, TTS recovery consistently showed the lowest accuracy among the four zero-shot tasks, although prediction accuracy remained above the expected baseline (Supporting Figure S6, Supporting Dataset S3). Notably, recovery accuracy was inversely associated with stop codon frequency, with TAA exhibiting the highest prediction accuracy despite being the least frequent stop codon (Supporting Figure S6). We further observed a significant negative correlation between genome-wide GC content and TTS recovery accuracy (Spearman’s ρ = −0.45, q = 2.4 × 10⁻□), with AT-rich genomes generally exhibiting higher accuracy. A similar trend was observed in angiosperms using PlantCAD2 (Supporting Figure S7, Supporting Dataset S4), suggesting that this relationship is not specific to BryoCAD but may reflect the influence of nucleotide composition on DNA language model predictions.

### BryoCAD captures sequence constraints associated with deleterious variation

We next investigated whether BryoCAD could predict the functional effects of genetic variants across 135 Marchantia genomes^18^. We first assessed the relationship between BryoCAD-predicted deleteriousness and minor allele frequency (MAF), as deleterious variants are expected to occur at lower frequencies due to purifying selection^19,20^. Zero-shot variant-effect prediction using *in silico* mutagenesis^19,20^ assigned more deleterious scores to mutations affecting conserved functional sites, particularly start-loss and stop-gain variants (Figure 4A, Supporting Dataset S5). We further extended this approach to structural variants (SVs) and found that deletions disrupting coding sequences, particularly frameshift deletions, received more deleterious scores than those in noncoding regions (Figure 4B, Supporting Dataset S5). Consistently, SNPs from natural populations were ranked by their zero-shot scores and classified into four percentile-based groups, with the top 0.5% representing the most deleterious variants. Mean MAF generally decreased with increasing predicted deleteriousness across three SNP categories, with the lowest frequencies observed for missense variants (Figure 4C, Supporting Dataset S5). Moreover, variants prioritized as deleterious by BryoCAD exhibited lower mean MAFs than those prioritized by PlantCAD2 across all percentile groups, suggesting that BryoCAD better captures sequence constraints associated with deleterious mutations. Together, these results demonstrate the potential of BryoCAD to assess the functional consequences of diverse genetic variants in bryophytes.

**Figure 4.**
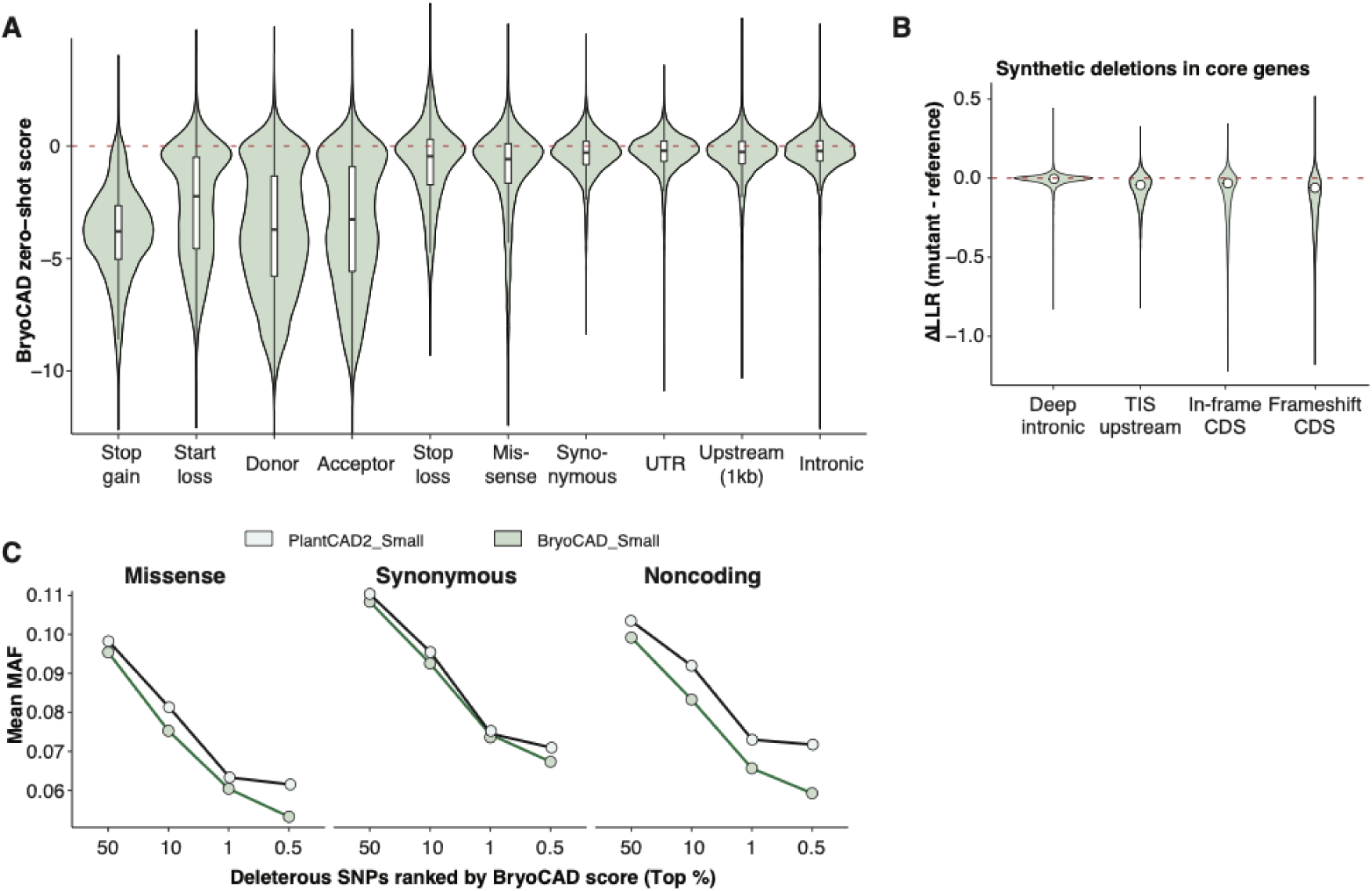
Zero-shot variant-effect prediction identifies putatively deleterious SNPs across 135 *Marchantia* genomes. (A) Violin plots showing the distribution of zero-shot variant-effect scores for SNPs generated by in silico mutagenesis. Predicted loss-of-function and splice-site variants exhibit scores consistent with stronger deleterious effects. (B) Violin plots comparing the distributions of zero-shot variant-effect scores across different functional classes of structure variants generated by in silico mutagenesis. (A) Minor allele frequency (MAF) distribution of SNPs prioritized as putatively deleterious by BryoCAD using zero-shot variant-effect prediction across 135 *Marchantia* genomes.

### BryoCAD improves transcript-level gene structure annotation in diverse bryophytes

To improve genome annotation accuracy, we adapted the GeneCAD framework^17^ by integrating BryoCAD with a fine-tuned gene prediction head, enabling accurate gene structure prediction directly from DNA sequences. BryoCAD outperformed alternative tools across all evaluation metrics (Figure 5A; Supporting Dataset S6), achieving coding-transcript F1 scores ranging from 0.53 to 0.63 and CDS-level F1 scores ranging from 0.84 to 0.89, demonstrating robust performance across diverse bryophyte genomes. We further assessed the effect of training-set diversity on gene annotation accuracy by comparing BryoCAD models trained on two or four bryophyte species. Although both models achieved high F1 scores across test species, the two-species model exhibited lower recall (Figure 5A, Supporting Dataset S6).

**Figure 5.**
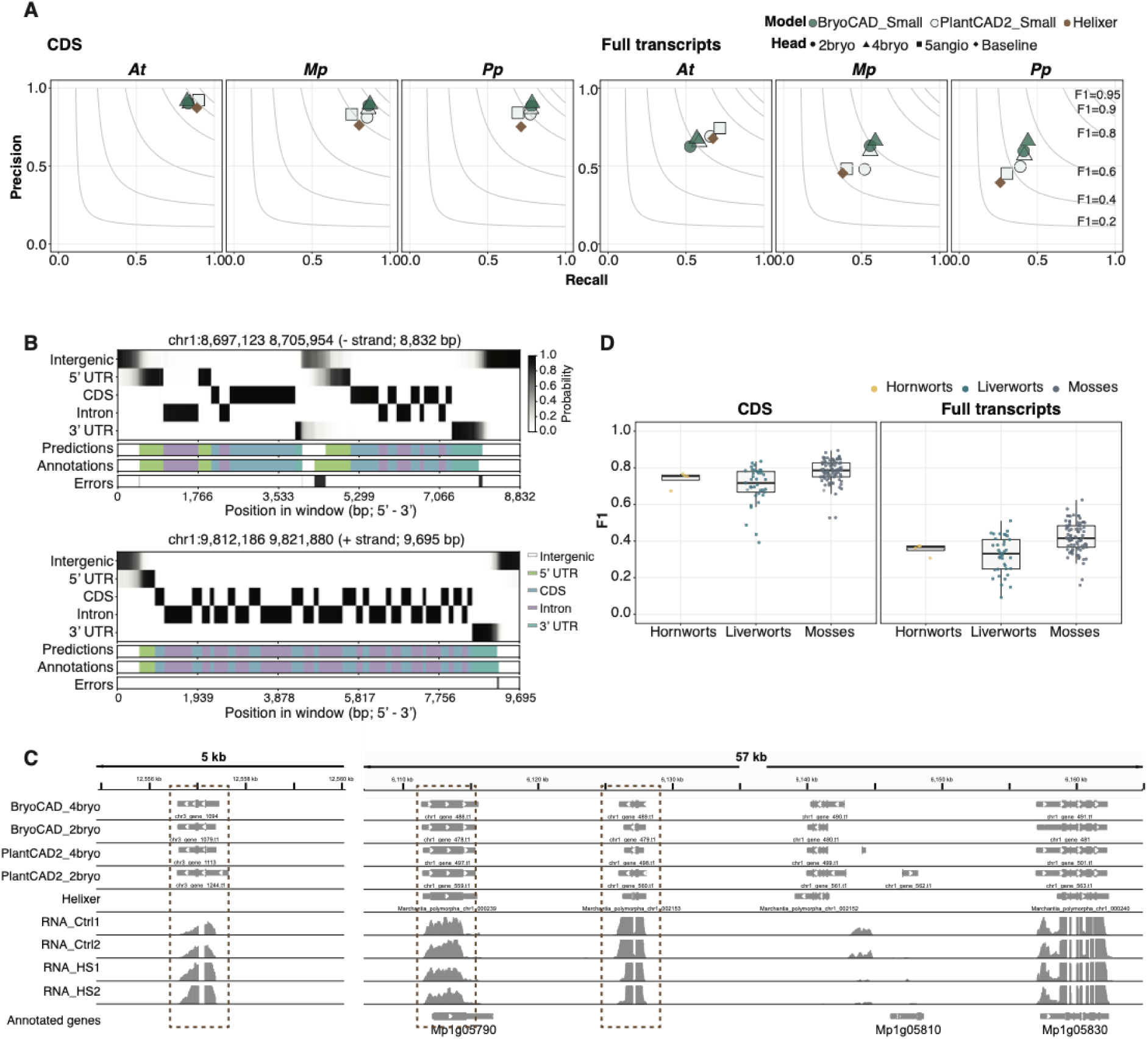
BryoCAD improves gene structure prediction and functional annotation in bryophytes. (A) Precision–recall plots at the transcript and CDS levels comparing PlantCAD2-5angio, BryoCAD-2bryo, BryoCAD-4bryo, and Helixer in liverwort, moss, and Arabidopsis genomes. (B) Representative gene-boundary probability profiles showing sharp peaks at translation initiation and termination codons and splice junctions, whereas 5′ and 3′ UTRs exhibit smoother probability gradients. The error track indicates discrepancies between model predictions and reference annotations. (C) Representative Integrative Genomics Viewer (IGV) views comparing predicted gene structures with reference annotations in Marchantia polymorpha. (D) Precision–recall plots at the transcript and CDS levels comparing model performance across held-out bryophyte species.

Boundary profiles showed sharper, higher-confidence transitions at translation initiation codons and splice junctions, whereas predictions across 5′ and 3′ UTRs exhibited smoother gradients (Figure 5B, Supporting Dataset S6). Visual inspection of representative genomic regions further revealed previously unannotated gene-like sequences in intergenic regions, as well as UTRs absent from the reference annotations (Figure 5C, Supporting Dataset S6). Among 1,116 candidate gene regions identified in the Marchantia polymorpha genome, approximately 300 were supported by multiple RNA-seq datasets. Notably, 13–40% of these candidate regions showed significant transcriptional responses to heat stress or other abiotic stresses, suggesting their potential involvement in environmental stress responses (Figure 5C; Supporting Dataset S6).

In addition, we extended gene annotation prediction to 123 bryophyte genomes (Figure 5D, Supporting Dataset S6). Across the three major bryophyte lineages, median CDS-level F1 scores ranged from 0.72 to 0.79, while coding-transcript F1 scores ranged from 0.33 to 0.42 (Figure 5D, Supporting Dataset S6). Together, these results demonstrate that BryoCAD supports genome-wide gene annotation across evolutionarily diverse bryophytes, while suggesting that expanding the diversity of training species may improve cross-species generalization, particularly by enhancing the recovery of annotated genes.

### Bryophyte-specific continued pre-training improves sequence prediction beyond land plants, particularly in algae

To further evaluate the cross-lineage generalization of BryoCAD, we assessed its zero-shot performance across four gene-boundary motif recovery tasks in evolutionarily divergent plant and algal genomes. Given that streptophyte algae are the closest algal relatives of land plants, we focused particularly on this lineage by including all publicly available genomes^21^. Compared with PlantCAD2, BryoCAD increased mean motif recovery accuracy from 26.3% to 37.5% across six algal species, while exhibiting lineage-dependent performance in lycophytes, ferns, and gymnosperms (Figure 6, Supporting Figure S8, Supporting Dataset S7). These results suggest that evolutionarily focused adaptation can enhance the predictive performance and cross-lineage transferability of DNA foundation models, extending their utility beyond the lineages used for pre-training.

**Figure 6.**
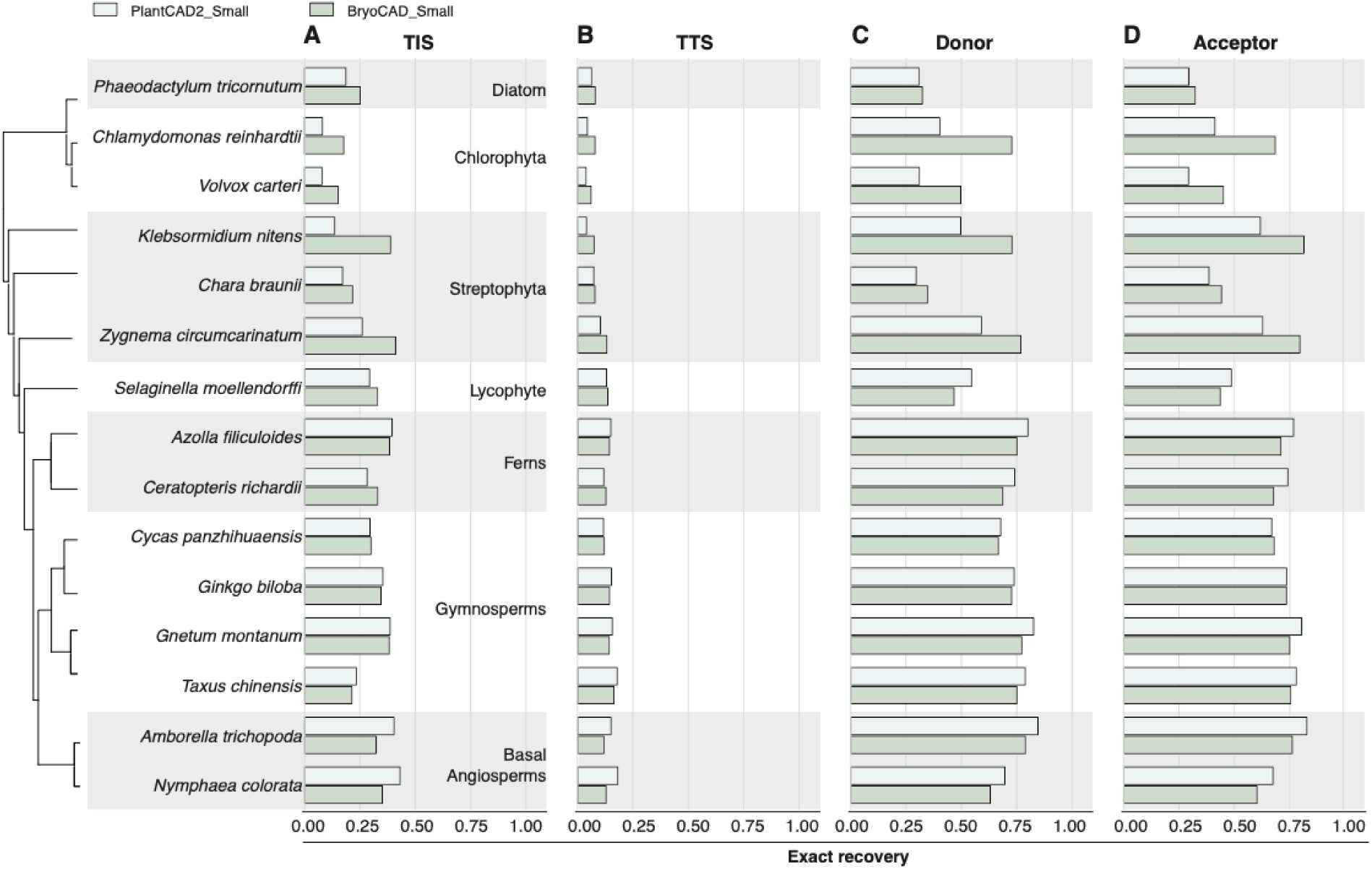
BryoCAD generalizes zero-shot gene-boundary motif recovery to divergent plant and algal lineages. (A–D) Evaluation of zero-shot motif recovery accuracy across 18 species representing diverse lineages of plants and algae. Bar plots show prediction accuracy for four genomic boundary tasks: (A) translation initiation sites (TISs), (B) translation termination sites (TTSs), (C) splice donor sites, and (D) splice acceptor sites.

## Discussion

Bryophytes are among the earliest land plants and occupy a crucial phylogenetic position for understanding plant terrestrialization. They represent the second-largest group of extant land plants, after flowering plants, and are widely distributed across diverse habitats^1,2^. They exhibit diverse adaptive strategies across environments and play important ecological roles in maintaining forest biodiversity, water retention, and nutrient cycling^22,23^. Although growing genomic and transcriptomic evidence suggests that bryophytes possess genomic and gene regulatory features distinct from those of flowering plants, analytical tools specifically designed for these lineages remain underdeveloped and rare, despite their evolutionary and ecological importance.

In this study, we present BryoCAD, a DNA LM tailored for bryophytes and a flexible framework that can be extended to other evolutionarily divergent and underrepresented plant lineages. Through continued pre-training of PlantCAD2, we demonstrate that BryoCAD improves performance across multiple genomic prediction tasks, including gene-boundary motif recovery, variant-effect prediction, chromatin accessibility prediction, and genome annotation, using both zero-shot and fine-tuning approaches. Notably, continued pre-training was performed on a single NVIDIA A100 GPU with 80 GB of memory, requiring approximately 317 GPU-hours for BryoCAD-Small and 1,123 GPU-hours for BryoCAD-Large, corresponding to approximately 13 and 47 days, respectively. In contrast, pre-training PlantCAD2-Large from scratch required approximately 43,000 H100 GPU-hours, while Evo2-40B required over 4 million H100 GPU-hours. These comparisons highlight the computational efficiency of continued pre-training for adapting existing DNA foundation models to evolutionarily divergent lineages. This highlighted the computational efficiency and accessibility of this approach for laboratories with limited computational resources.

BryoCAD’s superior performance in non-vascular plants compared with existing DNA foundation models, such as Evo2^12^ and NTv3^10^, highlights the importance of phylogenetic representation rather than simply increasing the number of training genomes when developing lineage-specific models^24^. This is consistent with the higher predictive performance of PlantCAD2 in angiosperms. However, the trade-off between lineage specialization and retention of predictive performance in the original training domain should also be considered, as its extent may vary across lineages and prediction tasks. Stop-codon recovery remained substantially lower than splice-site recovery. Stop-codon predictions also favored TAA across sites annotated as TAA, TAG, or TGA, even when TAA was not the predominant reference codon. This preference highlights the importance of distinguishing reference-codon usage, predicted codon frequencies, and exact recovery accuracy when assessing sequence prediction performance. The observed association with genome-wide GC content suggests a contribution from nucleotide composition, although the underlying mechanism remains unclear. Given that sequences surrounding TTSs can evolve rapidly and may contribute to phenotypic complexity in eukaryotes^25^, including plants^26–28^, the substantial differences in TTS recovery accuracy between bryophytes and angiosperms may reflect divergent sequence features, although their functional significance remains to be determined.

Furthermore, bryophyte genomes may retain more of the ancestral genomic architecture of early embryophytes: they have generally not undergone the extensive transposable element–driven genome expansion characteristic of many angiosperms^3^. Sequence features at gene boundaries learned from bryophytes may therefore lie closer to those of algal genomes than features learned predominantly from angiosperms. Consistent with this interpretation, BryoCAD’s improvements were most pronounced in algal lineages, including chlorophytes and a diatom outside Streptophyta, whereas its performance in vascular plants was comparable to or slightly below that of PlantCAD2. Together, our findings highlight the potential of phylogenetically focused pre-training to improve cross-lineage generalization and genomic sequence interpretation in underrepresented plant groups.

Although we demonstrated that BryoCAD can predict chromatin accessibility across different conditions in Marchantia (Supporting Figure S9), its ability to generalize across stress conditions and species requires further evaluation due to the limited availability of functional genomic datasets in bryophytes. Likewise, although BryoCAD can accurately recover gene-boundary motifs in algal lineages, its ability to predict genome-wide gene annotations requires further evaluation through fine-tuning of the gene prediction head using species-specific training data^17^. Gene-structure prediction is also constrained by annotation quality, particularly for 5′ and 3′ UTRs, where missing reference features can make it difficult to distinguish incomplete annotations from prediction errors. Nevertheless, completing 5′ and 3′ UTR annotations in bryophyte genomes represents a particularly valuable application of BryoCAD. Another promising direction is to apply BryoCAD to predict gene expression, promoter activity, and UTR-mediated translational regulation, particularly in the context of stress responses and cell-type-specific regulation. Ultimately, as more diverse datasets become available for bryophytes, we envision that BryoCAD and its predecessor, PlantCAD2, could be integrated with these resources as key components of predictive genomics frameworks across diverse plant lineages.

## Material & Methods

### Genome resources and continued pretraining

We assembled a collection of 126 bryophyte genome assemblies and associated annotations spanning hornworts, liverworts, and mosses. This collection comprised assemblies from 123 species included in the bryophyte pangenome resource of Dong *et al*. (2025)^3^, supplemented with reference assemblies of the three model organisms *Anthoceros agrestis*, *Marchantia polymorpha*, and *Physcomitrium patens*. Assembly names, taxonomic assignments, annotation versions, sources, and dataset roles are provided in Supporting Dataset S8. Seventeen assemblies, representing three hornworts, four liverworts, and ten mosses, were selected for continued pretraining. Selection prioritized broad representation of bryophyte orders, relatively compact genomes, and more complete gene annotations. Genome sequences and gene annotations were used as provided by the source databases, with preprocessing limited to the length, boundary, and ambiguous-base filters described below.

For the 17 selected assemblies, genomic sequences and gene coordinates were derived from the reference assemblies and annotations. Chromosomes or contigs shorter than 20 kb were excluded. Each annotated gene was extended by 5 kb on both sides to define a training region. Regions extending beyond a contig boundary, exceeding 50 kb including flanks, or containing more than 10% N bases were discarded. Negative-strand regions were reverse-complemented to gene orientation. Eligible chromosomes/contigs were shuffled across these 17 assemblies and assigned to validation until approximately 20% of their total sequence length was reached; remaining contigs were assigned to training. This partition preceded gene-region filtering and was not stratified by species. Contig assignments were saved in a split manifest.

BryoCAD-Small and BryoCAD-Large were initialized from their corresponding pretrained PlantCAD2 models^15^ and further pretrained on the selected bryophyte genomes. The architecture of each PlantCAD2 backbone was retained, and all model parameters were updated during continued pretraining. Gene regions were pooled without explicit species-balancing weights. At each presentation, an 8,192-bp crop was sampled uniformly from the available starts in the region. For dynamic masked language modeling, we selected 15% of eligible nucleotide positions, excluding padding and N tokens from the prediction targets. Of the selected positions, 80% were replaced with a mask token, 10% with a uniformly sampled A/C/G/T nucleotide, and 10% remained unchanged. Loss was evaluated only at selected positions. Repeat annotations were not used to modify the loss.

Continued pretraining used a learning rate of 5 x 10^-5, a per-device batch size of four, and eight gradient-accumulation steps. Training was configured for up to 50,000 optimizer updates, with a weight decay of 0.01, 1,000 warm-up steps, bfloat16 precision, and a random seed of 42. Validation was performed every 2,000 updates using up to the first 5,000 validation records with dynamic cropping and masking, and the checkpoint with the lowest validation loss was selected. The same continued-pretraining settings were applied to BryoCAD-Small and BryoCAD-Large.

### Orthogroup classification and genome-exposure categories

Protein sequences were obtained by translating the annotated coding sequence of the longest transcript of each gene. Missing sequences and proteins shorter than 50 amino acids, containing more than 30% unknown residues (X), or beginning with 20 or more consecutive X residues were excluded before orthogroup inference. Orthogroups were inferred using OrthoFinder v2.5.4^29^ and classified according to their presence across the 126 input genomes. OrthoFinder was run with default settings. An orthogroup was considered present in a genome if it contained at least one gene from that genome, irrespective of gene copy number. Core orthogroups were defined as those present in more than 80% of genomes (101-126 genomes), accessory orthogroups as those present in 2-100 genomes, and genome-specific orthogroups as those restricted to a single genome. Individual genes were assigned to the category of their corresponding orthogroup. Gene and orthogroup counts were summarized separately for each genome assembly. Multiple assemblies of the same species were treated independently, and genome-specific orthogroups were defined at the assembly level. Each protein left unassigned by OrthoFinder was represented as a singleton orthogroup and included in genome-specific gene and orthogroup counts. Gene-category proportions were calculated relative to the total number of classified genes, including genes represented by singleton orthogroups. Per-assembly core-gene counts are provided with the 126 bryophyte genome records in Supporting Dataset S8.

The 126 assemblies were classified into three mutually exclusive categories according to their representation in BryoCAD continued pretraining. Seen genomes comprised the 17 training assemblies. Unseen genomes from seen orders comprised 61 assemblies excluded from continued pretraining but belonging to orders represented in the training set. Unseen genomes from unseen orders comprised 48 assemblies belonging to orders absent from that set. These categories describe exposure during BryoCAD continued pretraining; they do not establish absence from the original pretraining corpora of PlantCAD2 or other comparator models. Exposure during GeneCAD head training was tracked separately.

### Zero-shot gene-boundary motif recovery

We evaluated zero-shot recovery of translation initiation sites (TIS), translation termination sites (TTS), splice donor sites, and splice acceptor sites following the motif-recovery framework of PlantCAD2^15^. For each gene, we selected the transcript with the longest coding sequence. Start and stop codon positions were obtained from explicit annotations when available; otherwise, they were inferred from the first and last three bases of the CDS, assuming that the annotated CDS included the stop codon. Splice donor and acceptor sites were defined as the first and last two bases of each intron, respectively, in the direction of transcription. Introns were identified from gaps between consecutive exons, using CDS segments when exon annotations were unavailable. For masked language models, 8,192-bp genomic windows were extracted with the target motif at the center, and all nucleotides of the motif were masked before prediction.

All 8,192-bp windows were oriented in the direction of transcription, with the target start or stop codon or splice-site dinucleotide positioned centrally. Sites with missing reference sequences, invalid coordinates, or insufficient flanking sequence to construct a complete window were excluded. Reference motifs were classified as canonical when they matched ATG for translation initiation, TAA, TAG, or TGA for translation termination, GT for splice donors, or AG for splice acceptors. Both canonical and noncanonical motifs were retained. No additional filtering was applied based on ambiguous bases within otherwise valid windows, and sites shared by different genes were not deduplicated. The same sequence-extraction and filtering procedures were applied across all benchmark datasets.

Exact motif recovery was defined as the proportion of evaluated sites for which the entire predicted motif matched the annotated sequence. For masked language models, all nucleotides of the target motif were masked simultaneously, and the highest-scoring nucleotide among A, C, G, and T was selected independently at each masked position in a single forward pass. Input-site counts and canonical/noncanonical motif composition for each of the 22 benchmark species and each task are provided in Supporting Datasets S9 and S10.

The principal benchmark included *A. agrestis*, *M. polymorpha*, *P. patens*, *A. thaliana*, *S. lycopersicum*, *O. sativa*, and *Z. mays*. We compared PlantCAD2-Small, PlantCAD2-Large, BryoCAD-Small, BryoCAD-Large, and Evo2 models with 1, 7, and 20 billion parameters^12^. For Evo2, forward and reverse-complement directional contexts were evaluated separately using its autoregressive prediction framework. The higher aggregate exact-recovery rate was retained for each model, species, and task combination. This best-direction summary used the evaluation sites themselves, with one direction selected per species and task rather than per site. Evo2 used greedy autoregressive generation (top_k = 1), conditioning successive motif bases on previously generated bases. Forward prompts comprised the sequence before the motif; reverse prompts comprised the reverse complement of the sequence after the motif. Thus TIS/TTS prompts contained 4,094 forward or 4,095 reverse bases, and splice prompts contained 4,095 bases in either direction. Reverse predictions were reverse-complemented before comparison with the annotated motif.

Motif recovery was evaluated across 22 species using two overlapping comparisons. Seven model species were evaluated with all seven models described above. A broader comparison across 18 species was restricted to PlantCAD2-Small and BryoCAD-Small and included three bryophytes, six algal species, one lycophyte, two ferns, four gymnosperms, and two angiosperms (*Amborella trichopoda* and *Nymphaea colorata*). The three bryophytes were included in both comparisons. Genome information and motif-input counts are provided in Supporting Datasets S9 and S10, respectively.

To assess generalization across bryophyte training-exposure categories, we evaluated BryoCAD-Small across the 126 assemblies. Exact recovery was calculated separately for each assembly and motif task using all evaluated genes, core and accessory genes combined, and core genes alone. For combined gene subsets, correct predictions and evaluated sites were summed before calculating recovery. Assembly-level recovery rates were compared using one-sided Welch t-tests with the alternative hypothesis that recovery was higher in seen genomes. Seen genomes were compared with all unseen genomes pooled and, separately, with each unseen category. Benjamini–Hochberg correction was applied across motif tasks and gene subsets within two separate families: pooled-unseen comparisons and comparisons with the two unseen categories. Gene-category recovery rates were also summarized separately for core, accessory, and genome-specific genes.

To examine compositional contributions to stop-codon prediction preferences, sites with annotated TAA, TAG, or TGA were analyzed separately. Exact recovery for each reference codon was calculated using all evaluated sites carrying that codon as the denominator. Predictions were grouped as TAA, TAG, TGA, or other triplets. Conditional prediction frequencies were calculated within each assembly and reference-codon category and summarized with equal weight across assemblies. The same TTS analyses were applied to PlantCAD2 predictions across 132 angiosperm species (Supporting Dataset S11), using the sequence-extraction and motif-recovery procedures described above.

Associations between genome GC content and predicted stop-codon frequencies were assessed using Spearman rank correlations. Each predicted codon was evaluated both as a fraction of all predictions and as a fraction of canonical stop-codon predictions. Analyses were performed separately for each reference codon and after pooling reference codons, with additional analyses within major lineages. To account for differences in reference-codon usage, conditional frequencies were also averaged with equal weight across the three reference codons within each assembly. Reference stop-codon frequencies were analyzed against genome GC content separately. Assemblies with missing GC values were excluded from these correlations. Benjamini–Hochberg correction was applied separately to the overall, within-lineage, equal-reference-weight, and reference-usage correlation families. Linear fits shown in scatterplots were descriptive.

To assess exact recovery beyond that expected from reference and prediction frequencies, a fixed-marginal null model was evaluated within each assembly. For the three reference codons and four prediction classes, 2,000 random contingency tables were sampled while preserving the observed row and column totals, equivalent to permuting prediction labels among sites within an assembly. Observed exact recovery was compared with the resulting null distribution using a one-sided Monte Carlo P value, calculated as (1 + the number of null values at least as large as observed)/2,001. Assembly-level P values were adjusted using the Benjamini–Hochberg procedure. An overall comparison used the equally weighted mean recovery across assemblies for both the observed and null values. The random seed was 20261008. This null model preserves model prediction preferences and tests their association with the annotated reference codons.

### Natural variants and in silico perturbation analyses

Natural SNPs were obtained from a *M. polymorpha* dataset comprising 135 accessions associated with the pangenome resource^6^. SNP contexts were annotated against the MpTak v6 reference genome and v6.1r2 gene annotation. The data were downloaded from https://marchantia.info/variants/data/Marpol_135_accessions/ (Marpol_135_accessions.vcf.gz and the accompanying accession list). Following PlantCAD (Zhai et al., 2025), each SNP was scored using a 512-bp sequence window with the variant position masked. The log-likelihood ratio (LLR) was calculated from the model-assigned conditional probabilities of the alternate and reference alleles:

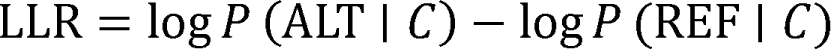

Here, ALT and REF denote the alternate and reference alleles, respectively, and C denotes the 512-bp sequence context with the variant position masked. More negative LLR scores indicate lower model compatibility of the alternate allele relative to the reference allele. Minor allele frequency (MAF) was calculated as the frequency of the less common allele among nonmissing genotype calls across the 135 accessions. SNPs were annotated by genomic context and orthogroup category using the reference genome annotation and orthogroup assignments.

To examine the relationship between variant scores and allele frequency, SNPs were grouped into synonymous coding variants, protein-altering variants (missense, start-loss, stop-gain, and stop-loss), and remaining variants. Within each group, SNPs were ranked by −LLR and assigned to four nonoverlapping percentile bands: 50th–90th, 90th–99th, 99th–99.5th, and 99.5th–100th percentiles. Mean MAF was calculated for each band to assess whether variants with lower model compatibility occurred at lower population frequencies.

Synthetic single-nucleotide variants were generated from *M. polymorpha* genes assigned to core orthogroups, using the MpTak v6 reference genome and v6.1r2 gene annotation. Each eligible unambiguous reference base was replaced with each of the other three nucleotides. Variants were assigned to coding-consequence, splice-flanking, UTR, promoter, or residual gene-body categories, and up to 10,000 variants per category were retained by reservoir sampling with seed 42. Strand-aware 1-kb upstream regions were used as putative promoters. Synthetic SNVs were scored using the same single-position masked LLR procedure as natural SNPs, following PlantCAD.

Synthetic structural variants were also evaluated in core-gene contexts. Four deletion classes were generated from annotated CDS-bearing transcripts: 100-bp deletions immediately upstream of the first CDS base; 9-bp frame-preserving CDS deletions; 10-bp frameshifting CDS deletions; and 10-bp deep-intron deletions. CDS deletions were at least 10 bp from segment boundaries, and deep-intron deletions were at least 100 bp from the adjacent CDS blocks. One eligible location per transcript and deletion class was selected where applicable, followed by random sampling of up to 1,000 candidates per class per genome (seed 42). Both reference and mutant sequences were 8,192 bp, oriented with transcription. Additional distal sequence compensated for deleted bases; boundary-truncated sequences and sequences with more than 5% N bases were excluded.

BryoCAD-Small evaluated 10 retained nucleotides on each side of the deletion. Corresponding retained bases were scored in the reference and mutant contexts. Each scored position was masked separately, and the log probability of its original base was averaged over valid A/C/G/T targets. The deletion score was the mutant minus reference mean masked log probability. Negative scores indicate reduced model compatibility of the retained junction-flanking bases after deletion. This localized pseudo-log-likelihood score is not the single-site SNP LLR and does not score deleted bases directly. Core-gene eligibility followed the orthogroup classification described above (Supporting Dataset S8). The generated deletion dataset comprised 12,000 events across 7,881 distinct gene loci: 4,000 events per species in *A. agrestis*, *M. polymorpha*, and *P. patens*, and 3,000 events per deletion category.

### Chromatin accessibility prediction using Low-rank Adaptation (LoRA)

We fine-tuned BryoCAD-Small for binary chromatin-accessibility prediction using LoRA^30^. Training data comprised control-condition ATAC-seq peaks from *M. polymorpha*^31^ (GSE290177), with sequences extracted from the MpTak v6 reference genome. Peak sets were generated locally from the source sequencing data. Positive examples were 512-bp windows centered on peak summits. Negative examples were non-overlapping 512-bp windows sampled across chromosomes and scaffolds containing annotated peaks, excluding any window overlapping a peak. Windows extending beyond sequence boundaries or containing ambiguous bases (N) were discarded. All remaining negative windows were included, without matching their GC content to positive examples or subsampling to balance the classes.

Data were partitioned by chromosome to separate training and validation loci. Chromosomes 7 and 8 were reserved for validation, and the remaining eligible chromosomes and scaffolds were used for training. The resulting datasets contained 32,186 positive and 265,052 negative training examples, and 8,858 positive and 62,228 negative validation examples.

LoRA adapters were applied to the x_proj, in_proj, and out_proj modules with rank 8, alpha 32, and dropout 0.1. Fine-tuning used AdamW with an initial learning rate of 1 x 10^-3, a linear learning-rate schedule with 50 warmup steps, and unweighted cross-entropy loss. Models were trained for 4,000 optimizer steps using a per-device batch size of four and 16 gradient-accumulation steps, corresponding to an effective batch size of 64 per device. The random seed was set to 42, and the final checkpoint was used for evaluation.

Transfer performance was evaluated without further fine-tuning using *M. polymorpha* heat-stress ATAC-seq data from GSE290177^31^ and *A. thaliana* control and heat-stress DNase-seq data from GSE145742^32^. M. polymorpha heat-stress evaluation was restricted to chromosomes 7 and 8, which were excluded from model training and also used for control-condition validation. Test examples were prepared using the same window-generation, negative-sampling, and sequence-filtering procedures as the training data. Performance was assessed using AUROC calculated from positive-class probabilities and F1 calculated from the class with the higher predicted probability, without decision-threshold optimization.

### GeneCAD head training and gene-structure prediction

We trained gene-annotation heads using GeneCAD v0.4.0^17^, with BryoCAD-Small or PlantCAD2-Small as the pretrained backbone. Two bryophyte training sets were used: 2bryo, comprising *M. polymorpha* and *A. agrestis*, and 4bryo, which additionally included *Ceratodon purpureus* and *Sphagnum magellanicum*. Heads were trained with each backbone on both datasets. An additional PlantCAD2-Small head was trained using the five-angiosperm training set defined in GeneCAD (*A. thaliana, Glycine max, Hordeum vulgare, O. sativa, and Populus trichocarpa*), providing a comparator trained on angiosperm annotations.

Reference annotations were restricted to protein-coding genes, retaining one representative transcript per gene based on the greatest summed CDS length. Ties were resolved by transcript genomic span and then transcript identifier. Exon, CDS, and untranslated-region annotations associated with the selected transcripts were retained.

The pretrained backbones were frozen during head training. Annotation heads used the GeneCAD legacy architecture, with a token embedding dimension of 128, a hidden dimension of 768, and eight attention layers. Training used 8,192-bp windows, an intergenic proportion of 0.15, and automatically calculated class weights. Following the GeneCAD plant-training defaults, windows within each chromosome or contig were randomly partitioned into approximately 95% training and 5% validation samples (seed 42). The species configurations enabled both training and validation for the species in each training set and excluded *P. patens* from both splits. Training was configured for five epochs with a batch size of eight and five gradient-accumulation steps, yielding an effective batch size of 40. A constant learning rate of 1 x 10^-4 and a weight decay of 1 x 10^-5 were used. For each configuration, the checkpoint with the highest overall validation entity-level F1 was used for prediction.

Gene prediction was evaluated across the complete assemblies of *A. thaliana, M. polymorpha,* and *P. patens*. For heads trained on the 2bryo and 4bryo datasets, *A. thaliana* and *P. patens* were absent from head training, whereas *M. polymorpha* was included. *A. thaliana* was included in the five-angiosperm head-training set. *P. patens* was represented in BryoCAD continued pretraining.

Predictions were compared with reference annotations using a modified version of the GeneCAD evaluation script. Base-level performance was calculated separately for CDSs, 5’ UTRs, 3’ UTRs, and the combined exonic regions. For each region type, annotated intervals were merged within each sequence and strand to avoid counting overlapping bases more than once. True-positive bases were those shared by the predicted and reference intervals; predicted-only and reference-only bases were counted as false positives and false negatives, respectively. Exonic and UTR comparisons were reported only when the reference annotations supported these features.

Structural agreement was evaluated separately from base-level performance. CDS-segment matches required identical sequence identifiers, strands, and start and end coordinates. CDS transcript matches required exact agreement of the complete CDS-segment structure, irrespective of UTR boundaries. Transcript and locus comparisons used maximum one-to-one matching, with a CDS-locus match requiring at least one exactly matching CDS transcript structure. Precision, recall, and F1 were calculated as TP/(TP + FP), TP/(TP + FN), and 2TP/(2TP + FP + FN), respectively. Metrics with undefined denominators were reported as unavailable.

Publicly available *M. polymorpha* RNA-seq datasets were used to assess transcriptional support and stress responsiveness of GeneCAD-predicted genes. GSE290176 provided Tak-1 samples subjected to heat stress or abscisic acid (ABA) treatment and their corresponding controls (PMID: 41391150), whereas PRJEB48734 provided samples subjected to cold, heat, salt, mannitol, nitrogen deficiency, high light, or darkness, together with two control groups (PMID: 36813788). Paired-end reads were aligned to the *M. polymorpha* MpTak_v6.1r2 reference genome using HISAT2 v2.2.1. Paired-end fragments were quantified across all GeneCAD-predicted genes using featureCounts (Subread v2.0.6) in unstranded mode, with CDS features grouped by transcript and transcript identifiers mapped to gene identifiers.

Novel annotation candidates were defined as GeneCAD-predicted loci whose complete gene spans did not overlap any reference gene interval, irrespective of strand and without additional flanking-distance exclusion. Each candidate was represented by one counted transcript. Expressed candidates required raw fragment counts >10 in at least two individual samples within the corresponding experiment, considering treatment and control samples together. These candidates were regarded as previously unannotated predicted loci rather than experimentally validated genes.

Differential expression was analyzed independently for each treatment–control comparison using PyDESeq2 v0.5.4. All predicted transcripts with raw counts greater than 10 in at least two samples per comparison were included in normalization, model fitting, and testing, after which results for novel candidates were extracted. Genes were considered differentially expressed if their Benjamini–Hochberg-adjusted P values were less than 0.05 and their absolute unshrunk log2 fold changes were at least 1. For PRJEB48734, candidates were required to meet these criteria against both control groups with concordant directions.

## Data availability

Source code and analysis workflows are available at https://github.com/yenmr/BryoCAD. Pretrained model weights are available at https://huggingface.co/bryocad.

## Acknowledgment

We thank the Academia Sinica Grid Computing Centre (ASGC; AS-CF11-114-A11) and the Texas Advanced Computing Center at The University of Texas at Austin (#MCB24097) for providing computational resources. This study was supported by Academia Sinica (AS-CDA-113-L03 to TYW) and the National Science Foundation (NSF; #2532633 to JZ and SKH).

## Author’s contribution

TYW conceived the project. TYW, SKH, JZ, and MRY designed the study. MRY and JZ performed the analyses. TYW and MRY wrote the manuscript with input from all authors. TYW, SKH, JZ, and MRY reviewed and edited the manuscript.

## Supporting Figures

**Supporting Dataset S1.** Prediction results from all models across seven benchmark species. Related to Figure 2.

**Supporting Dataset S2.** Prediction results across 126 bryophyte species. Related to Figure 3.

**Supporting Dataset S3.** Prediction results of TTS across 126 bryophyte species. Related to Supporting Figure S6.

**Supporting Dataset S4.** Prediction results of TTS across 123 angiosperm species. Related to Supporting Figure S7.

**Supporting Dataset S5.** Prediction results across 135 Marchantia species. Related to Figure 4.

**Supporting Dataset S6.** Genome-wide gene structure prediction results across 126 bryophyte species and their expressions under stress conditions. Related to Figure 5.

**Supporting Dataset S7.** Prediction results across algal and other land plant lineages. Related to Figure 6.

**Supporting Dataset S8.** Genome information for 126 bryophyte assemblies, including continued-pretraining status and core-gene counts.

**Supporting Dataset S9.** Genome information for 22 distinct species used in the seven-species model benchmark and the 18-species evolutionary-lineage comparison. The comparisons share three bryophytes; the lineage comparison excludes *A. thaliana*, *S. lycopersicum*, *O. sativa*, and *Z. mays*. The table labels Model_Benchmark_7 and Transfer_15 distinguish the seven original benchmark species from the 15 additional species, rather than membership in the 18-species comparison.

**Supporting Dataset S10.** Motif counts for the 22 benchmark species by task, including retained input sites, canonical and noncanonical motif counts, and exclusions.

**Supporting Dataset S11.** Genome and annotation information for 132 angiosperm species used in the PlantCAD2 stop-codon recovery and GC-associated prediction-preference analyses.

---

**Supporting Figure S1.**
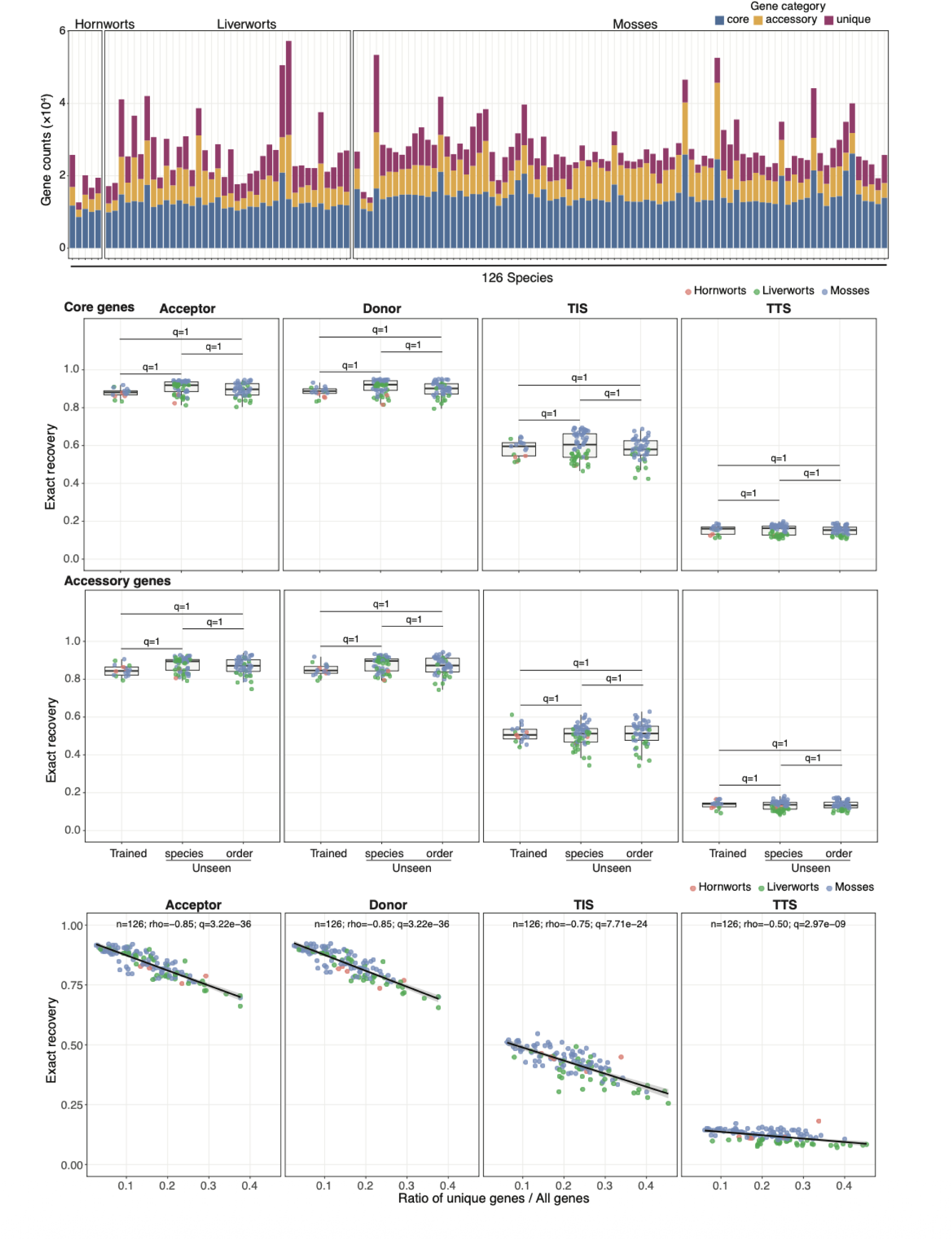
Gene set composition and motif recovery accuracy across bryophyte genomes. (A) Bar plots showing the numbers of core, accessory, and unique genes in each species. Core genes were defined as those present in more than 80% of genomes (101–126 genomes), accessory genes as those present in 2–100 genomes, and unique genes as those restricted to a single genome. (B) Box plots comparing exact motif recovery accuracy between core and accessory genes across three genome categories: seen genomes (n = 17), included in training; unseen species (n = 61), excluded from training but belonging to orders represented in training; and unseen orders (n = 48), absent from training. (C) Scatter plots showing the relationship between the proportion of unique genes and exact motif recovery accuracy for four genomic boundary motifs across species.

**Supporting Figure S2.**
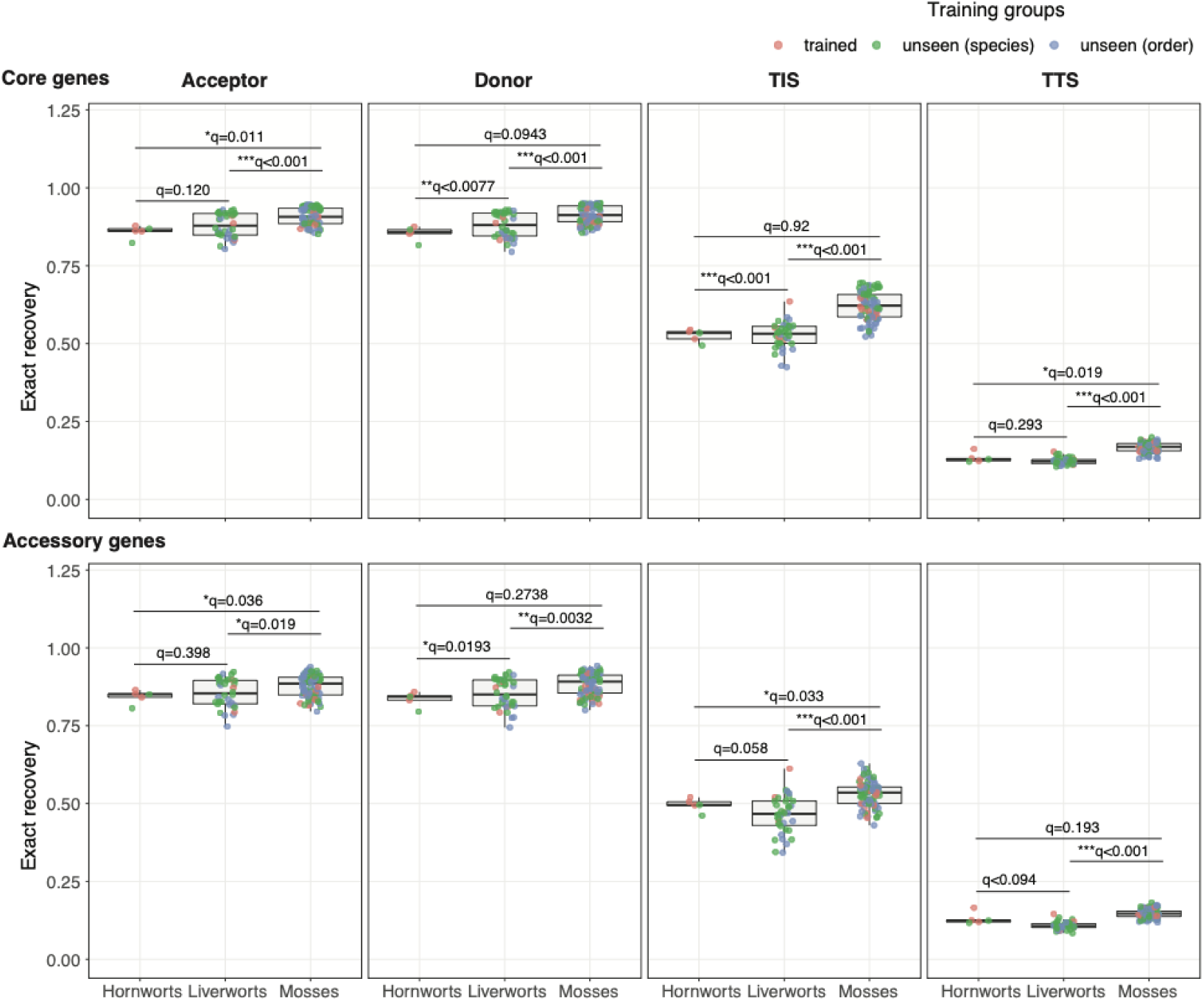
Comparison of motif recovery accuracy across three bryophyte lineages. (A and B) Box plots comparing exact recovery accuracy for four genomic boundary motifs in (A) core genes and (B) accessory genes across hornworts, liverworts, and mosses. In both panels, dots represent individual data points, center lines indicate medians, and boxes span the first and third quartiles. Statistical significance was assessed using one-sided Welch’s t-tests with Benjamini–Hochberg correction for multiple comparisons. Adjusted P-values are reported as q-values.

**Supporting Figure S3.**
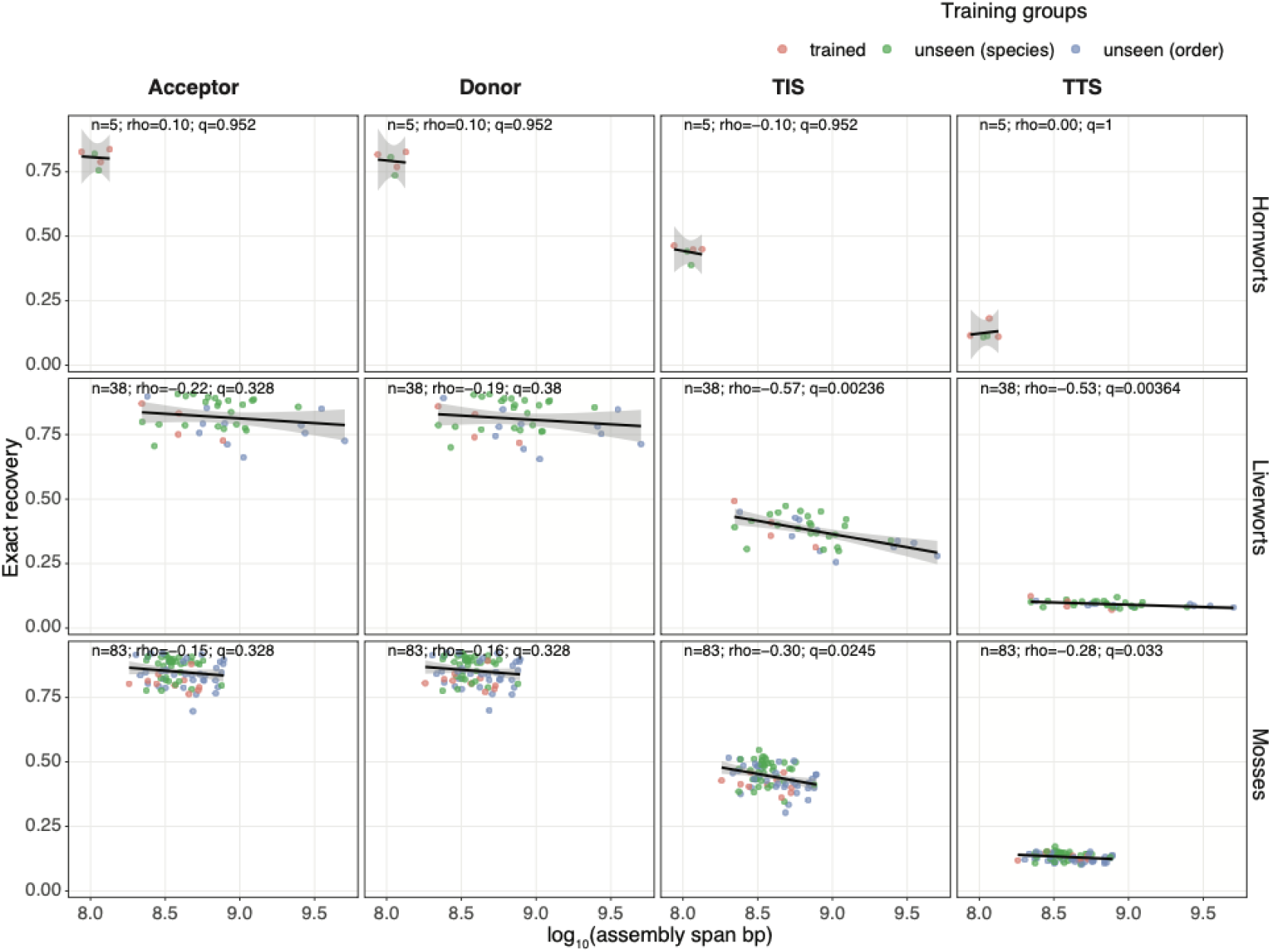
Relationship between genome size and motif recovery accuracy across bryophytes. Scatter plots showing the correlation between genome size and exact recovery accuracy for four genomic boundary motifs across species. Each dot represents one species. Spearman’s rank correlation coefficients (ρ) and Benjamini–Hochberg-adjusted *P* values are reported. Solid lines represent ordinary least-squares (OLS) regression fits, with gray shaded areas indicating 95% confidence intervals (CIs).

**Supporting Figure S4.**
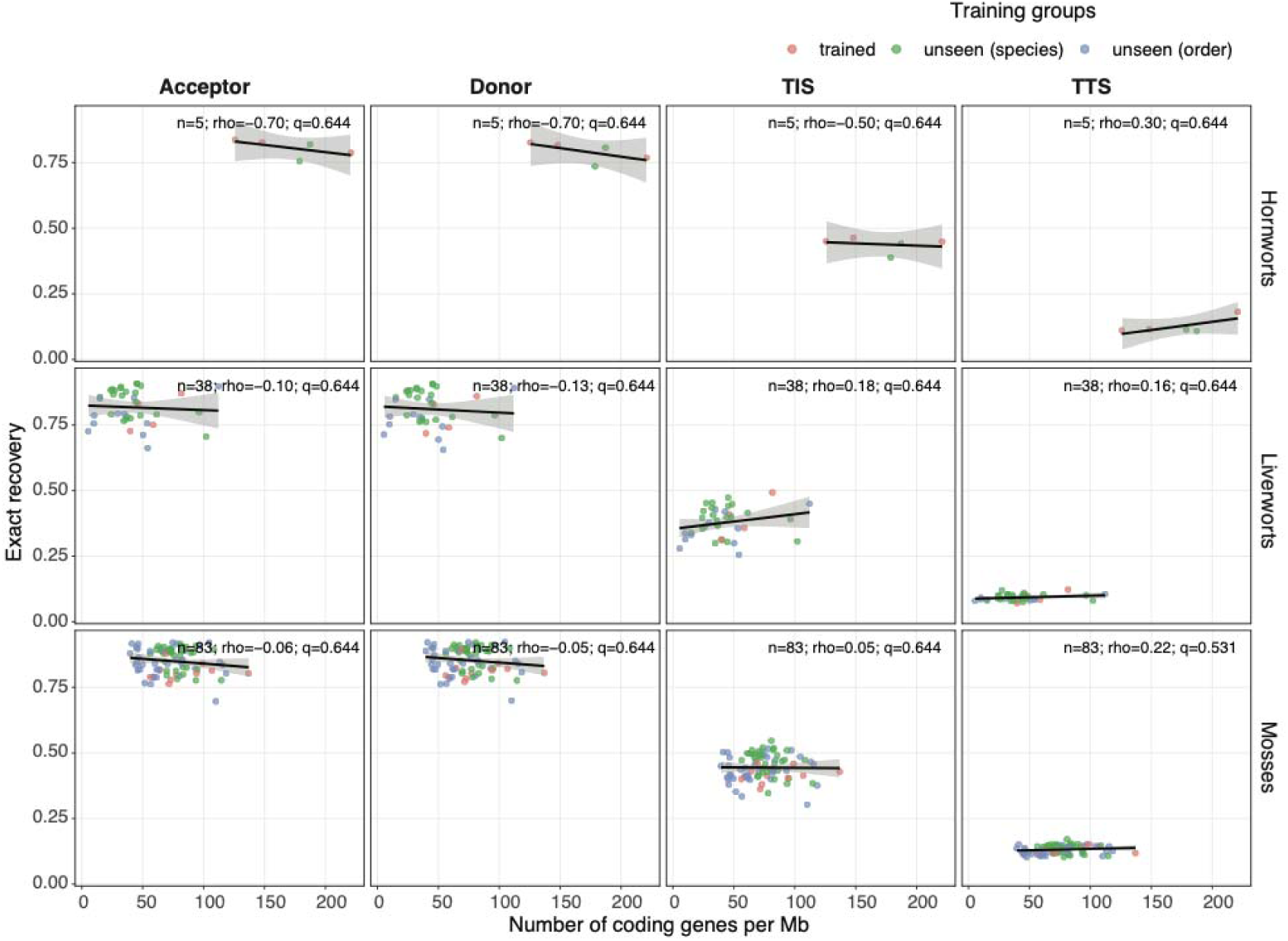
Relationship between gene density and motif recovery accuracy across bryophytes. Scatter plots showing the correlation between gene density and exact recovery accuracy for four genomic boundary motifs across species. Each dot represents one species. Spearman’s rank correlation coefficients (ρ) and Benjamini–Hochberg-adjusted P-values are reported. Solid lines represent ordinary least-squares (OLS) regression fits, with gray shaded areas indicating 95% confidence intervals (CIs).

**Supporting Figure S5.**
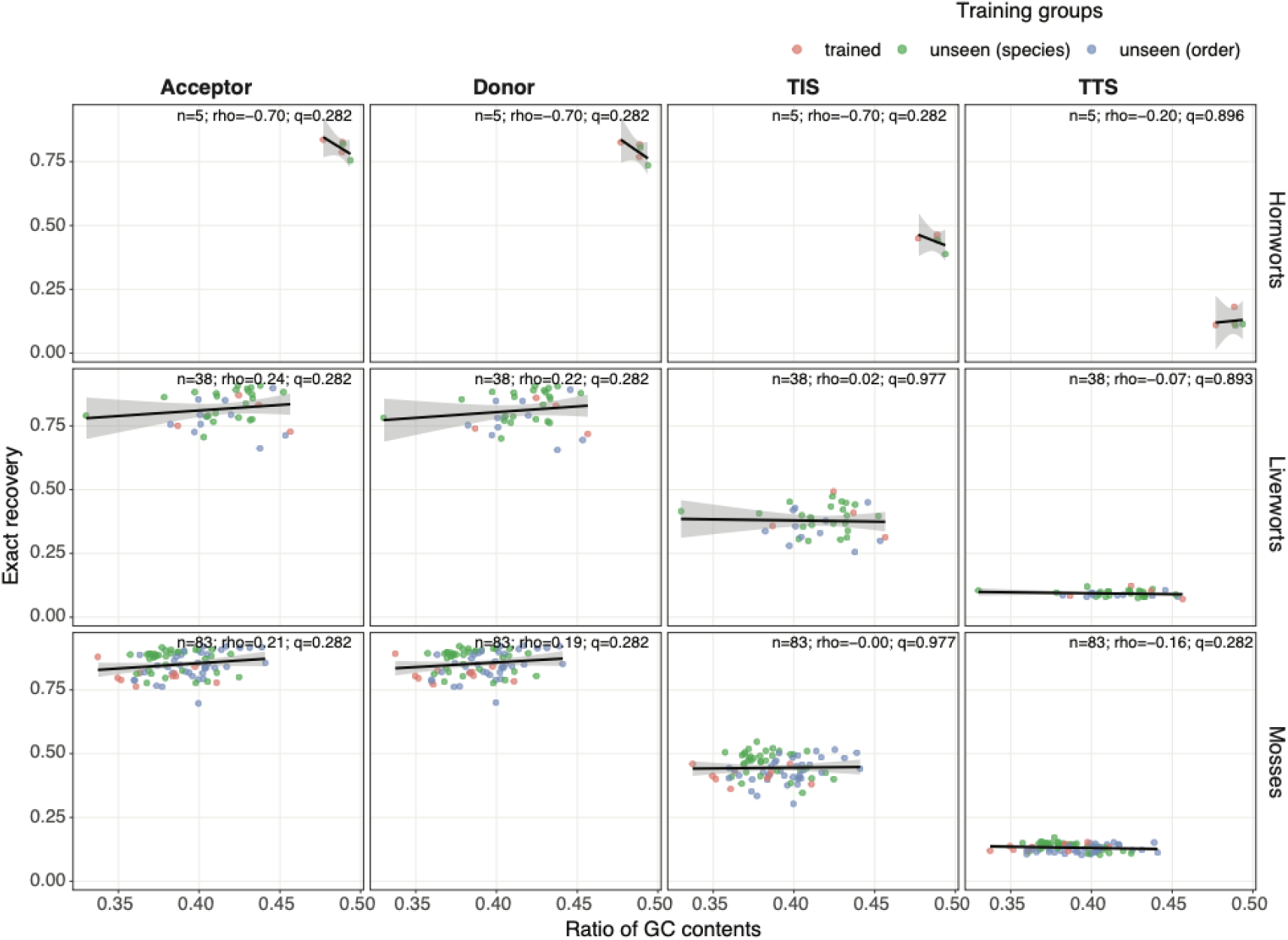
Relationship between GC content and motif recovery accuracy across bryophytes. Scatter plots showing the correlation between genomic GC content and exact recovery accuracy for four genomic boundary motifs across species. Each dot represents one species. Spearman’s rank correlation coefficients (ρ) and Benjamini–Hochberg-adjusted *P* values are reported. Solid lines represent ordinary least-squares (OLS) regression fits, with gray shaded areas indicating 95% confidence intervals (CIs).

**Supporting Figure S6.**
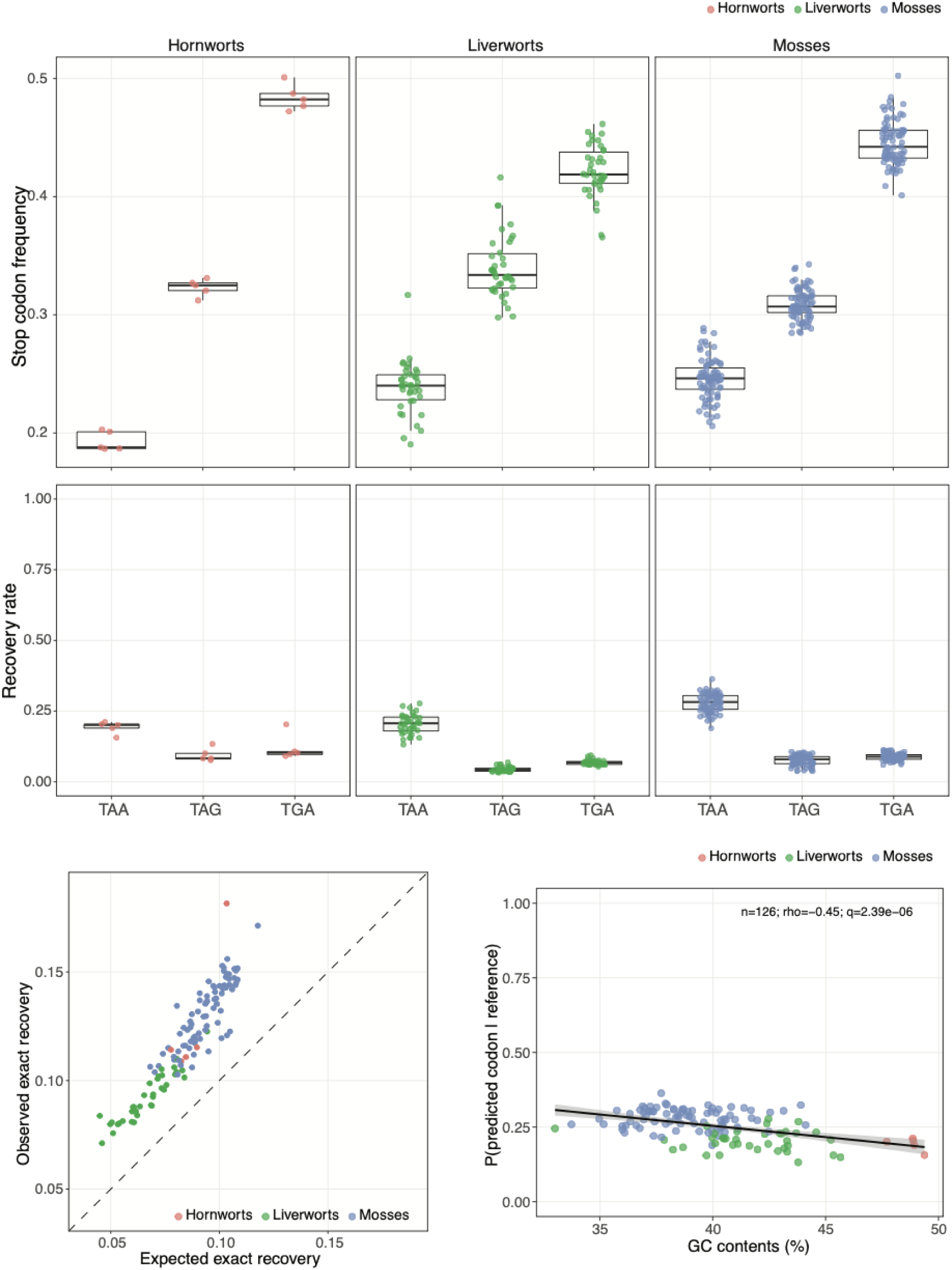
Stop codon composition and translation termination site recovery accuracy across bryophytes. (A) Distribution of stop codon frequencies across bryophyte lineages and their corresponding exact recovery accuracies. (B) Distribution of observed and expected exact recovery accuracies across species. The dashed line indicates the expected accuracy under random guessing. (C) Scatter plots showing the correlation between GC content and exact recovery accuracy across species. Each dot represents one species. Spearman’s rank correlation coefficients (ρ) and Benjamini–Hochberg-adjusted P-values are reported. Solid lines represent ordinary least-squares (OLS) regression fits, with gray shaded areas indicating 95% confidence intervals (CIs).

**Supporting Figure S7.**
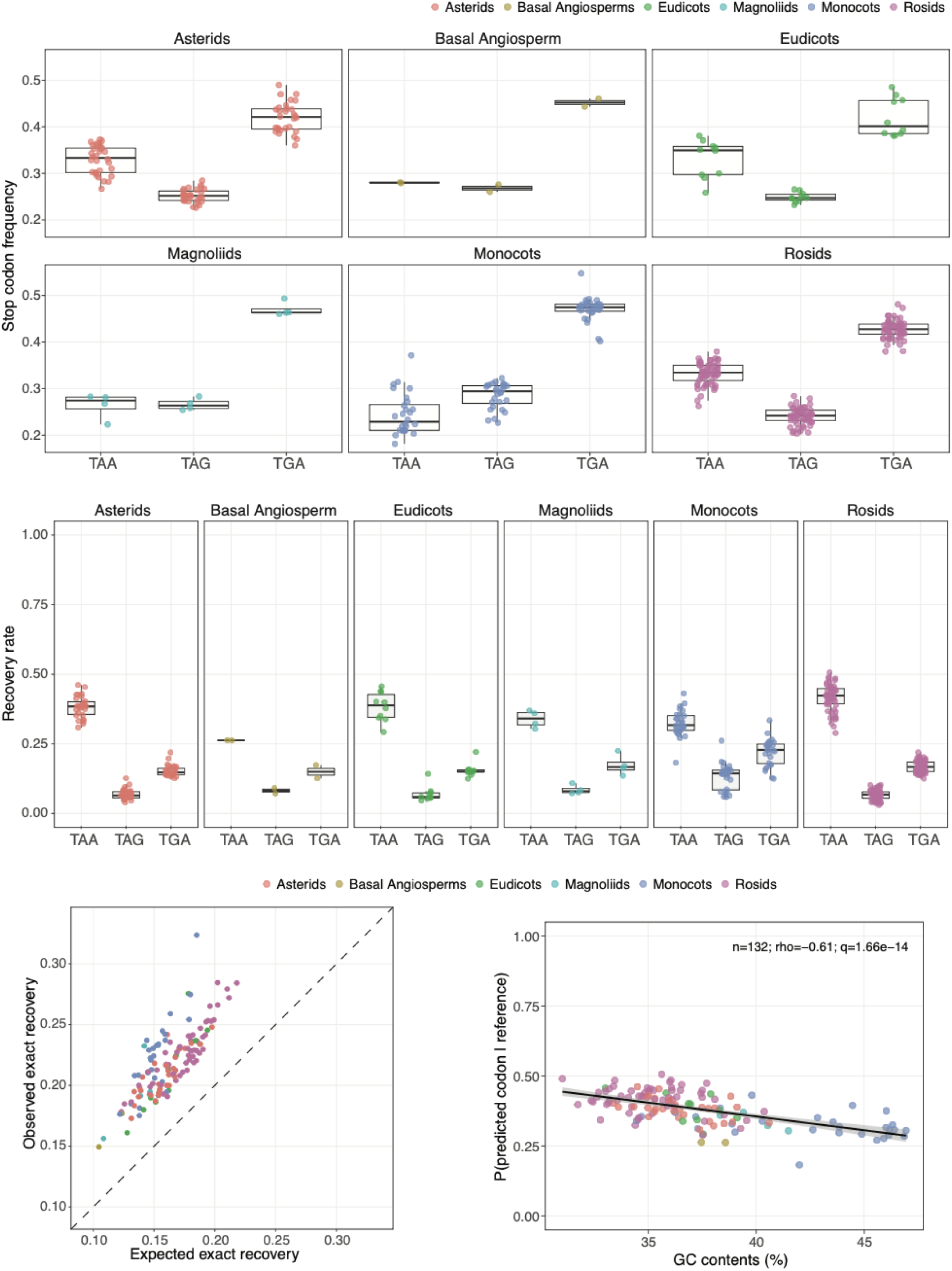
Stop codon composition and translation termination site recovery accuracy across angiosperms. Same analyses as in Supporting Figure S6, but performed on angiosperm genomes using PlantCAD2 for zero-shot motif recovery prediction.

**Supporting Figure S8.**
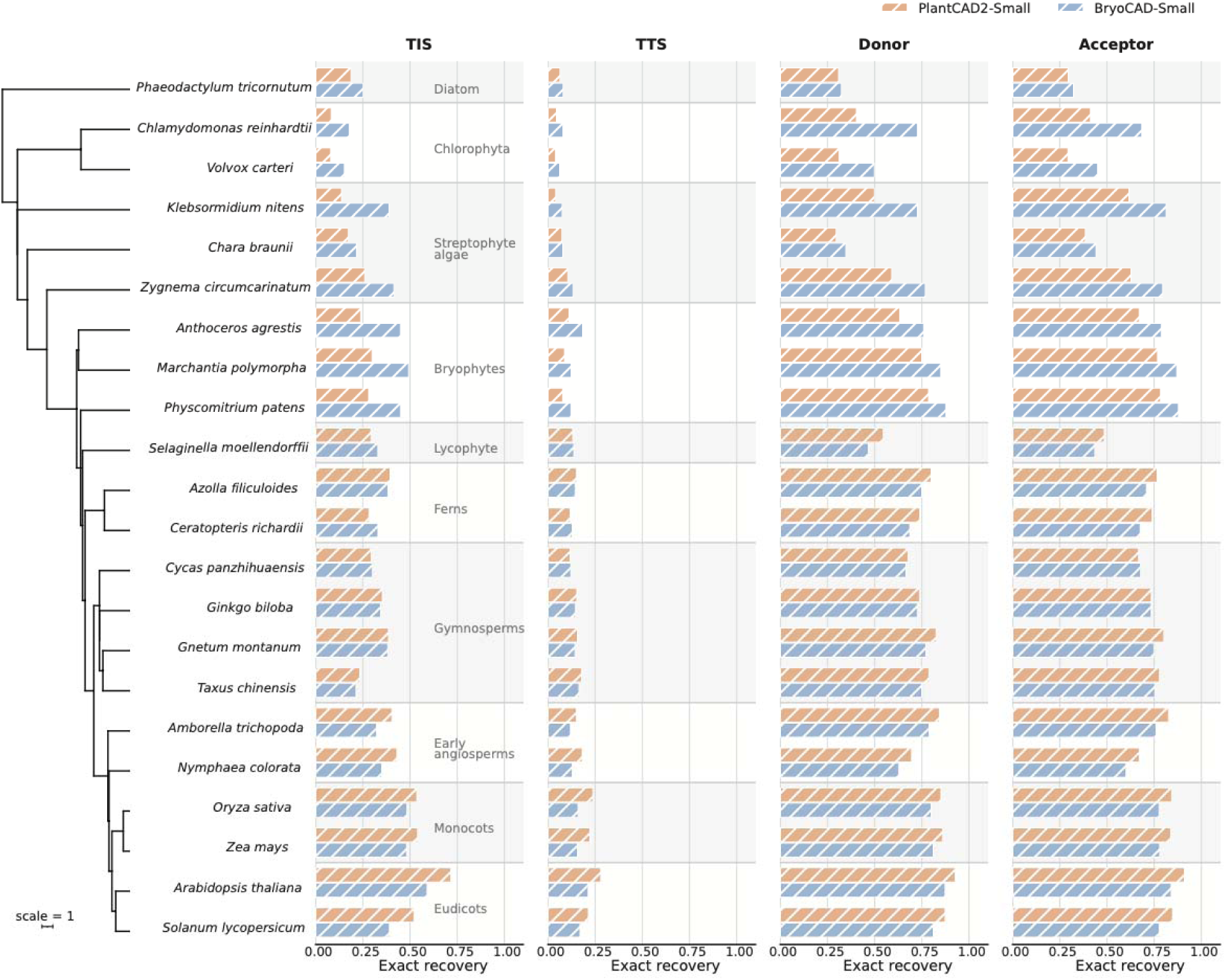
**BryoCAD generalizes zero-shot gene-boundary motif recovery to divergent plant and algal lineages**. Similar to Figure 6, but showing all 22 species and their phylogenetic relationships.

**Supporting Figure S9.**
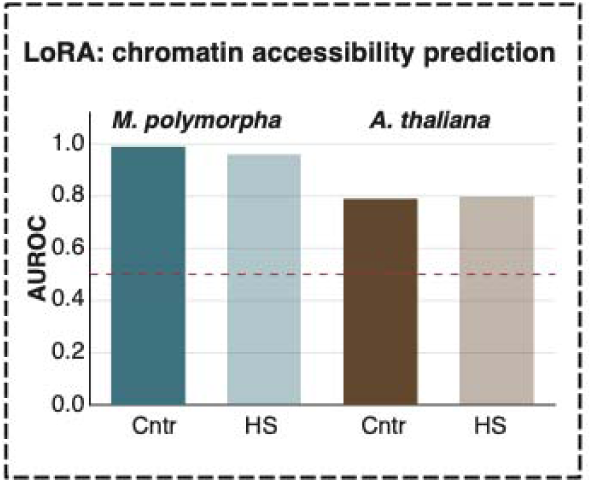
BryoCAD predicts chromatin accessibility in *Marchantia* and *Arabidopsis* using LoRA fine-tuning. Bar plots showing F1 scores for chromatin accessibility prediction in *Marchantia polymorpha* and *Arabidopsis thaliana* under control and heat stress conditions.

